# SARS-CoV-2 infecting the inner ear results in potential hearing damage at the early stage or prognosis of COVID-19 in rodents

**DOI:** 10.1101/2020.12.23.423942

**Authors:** Xia Xue, Yongan Tian, Mingsan Miao, Jianyao Wang, Wenxue Tang, Yaohe Wang, Jianbo Liu, Hongen Xu, Jinxin Miao

## Abstract

**Objectives:** In order to find out the association between the sensorineural hearing loss and COVID-19, we detected the expression ACE2 and TMPRSS2 in the mouse the hamster.

**Design:** Using the public data from NCBI and GISAID, we assessed the expression of ACE2 and TMPRSS2 at the transcriptomic, DNA, and protein levels of ACE2 in the brain, inner ear, and muscle from the golden Syrian hamster (*Mesocricetus auratus*) and mouse (*Mus musculus*).

**Results:** We identified ACE2 and TMPRSS2 expressed at different level in the inner ear and brain at DNA and transcriptomic levels of both mouse and the hamster. The protein expression shows a similar pattern of the brain and inner ear, while the expression of ACE2 from the inner ear was relatively higher than it from the muscle.

**Conclusion:** SARS-CoV-2 could infect the hearing system potentially and SSNHL could be a characteristic to detect asymptomatic patients of COVID-19.

## Introduction

Hundred thousand death caused by a coronavirus (SARS-CoV-2) disease 2019 (COVID-19) within a few months highlights the importance of early diagnosis of COVID-19. Meanwhile, the increasing trend of the confirmed cases draws more attention to the treatment and prognosis of COVID-19 patients. Some recent clinical studies observed some un-specific symptoms of COVID-19, including headache, diarrhea, and sudden sensorineural hearing loss (SSNHL) (Kilic et al. 2020; Harenberg, Jonas, and Trecca 2020); one case has reported profound sensorineural hearing damage in a 60-year-old COVID-19 patient (Degen, Lenarz, and Willenborg 2020). To our acknowledge, the ACE2 receptor is a key for SARS-Cov-2 entry human cells while TMPRSS2 is highly involved in virus replication, the co-expression of which two could mainly determine the level of damage to the hosts. It has been previously investigated that the expression of ACE2 and TMPRSS2 was high in the intestine and kidney of humans compared to the brain and lung. Although several clinical studies confirmed the most common symptoms of COVID-19, SSNHL were also found but not draw much attention due to they are similarity to the symptoms of the flu or cold. SARS-CoV-2 could infect many organs, including the hearing center, which could invade the cochlear nerve and lead to SSNHL and cause hearing damage after treatment of the patients with COVID-19. We have already demonstrated the golden hamster (*Mesocricetus auratus*) could be a better small animal model for COVID-19 compared to a mouse (*Mus musculus*) for understanding the efficiency and possibility of SARS-CoV-2 attacking human cells with certain or potential consequences.

## Methods

The procedures in this study involving animals were reviewed and approved by the Animal Experimentation Ethics Committee of the Henan University of Chinese Medicine (No. DWLL202001301). Raw RNA-seq data of transcriptomes were collected from NCBI. The coding sequence of ACE2 and TMPRSS2 was aligned with Clustral Omega (V1.2.3) and reconstructed phylogeny, calculated distance by using MEGA v5.2. 1000 maximum number of iterations were applied with Kimura two factor correction method for genetic distance prediction. Cufflinks v2.2.1 with Bowtie2 (Langmead and Salzberg 2012) were used for expression analysis of transcriptomes from different tissues in humans, mice, and hamsters. We next ran quantitative real-time PCR (qRT-PCR) and western blot on seven tissues from both mouse and the hamster to quantitate the expression of ACE2 and TMPRSS2. All experimental details are included in the supplementary file.

## Results

Gene *ACE2* and *TMPRSS2* of the mouse and hamster are over 80% similarity to humans (figure 1A). We found *ACE2* expressed higher in the heart of humans and slightly higher in the brain of both human and mouse, while *TMPRSS2* shows high expression in the lungs of both human and mouse but lower in the brain (figure 1B). In addition, we found a similar pattern of the hamster and human; Both *ACE2* and *TMPRSS2* were highly expressed in the spleen and relatively high in the liver, but not in the brain (figure 1C). The mRNA expression of *ACE2* shows high in the inner ear of both the mouse and hamster, while *TMPRSS2* was much higher expressed in the lung compared to the inner ear. However, both genes showed lower expression levels in the brains of these two animal models. The protein expression shows a similar pattern of the brain and inner ear, while the expression of *ACE2* from the inner ear was relatively higher than it from the muscle (figure 1D, E, F).

**Figure 1.**
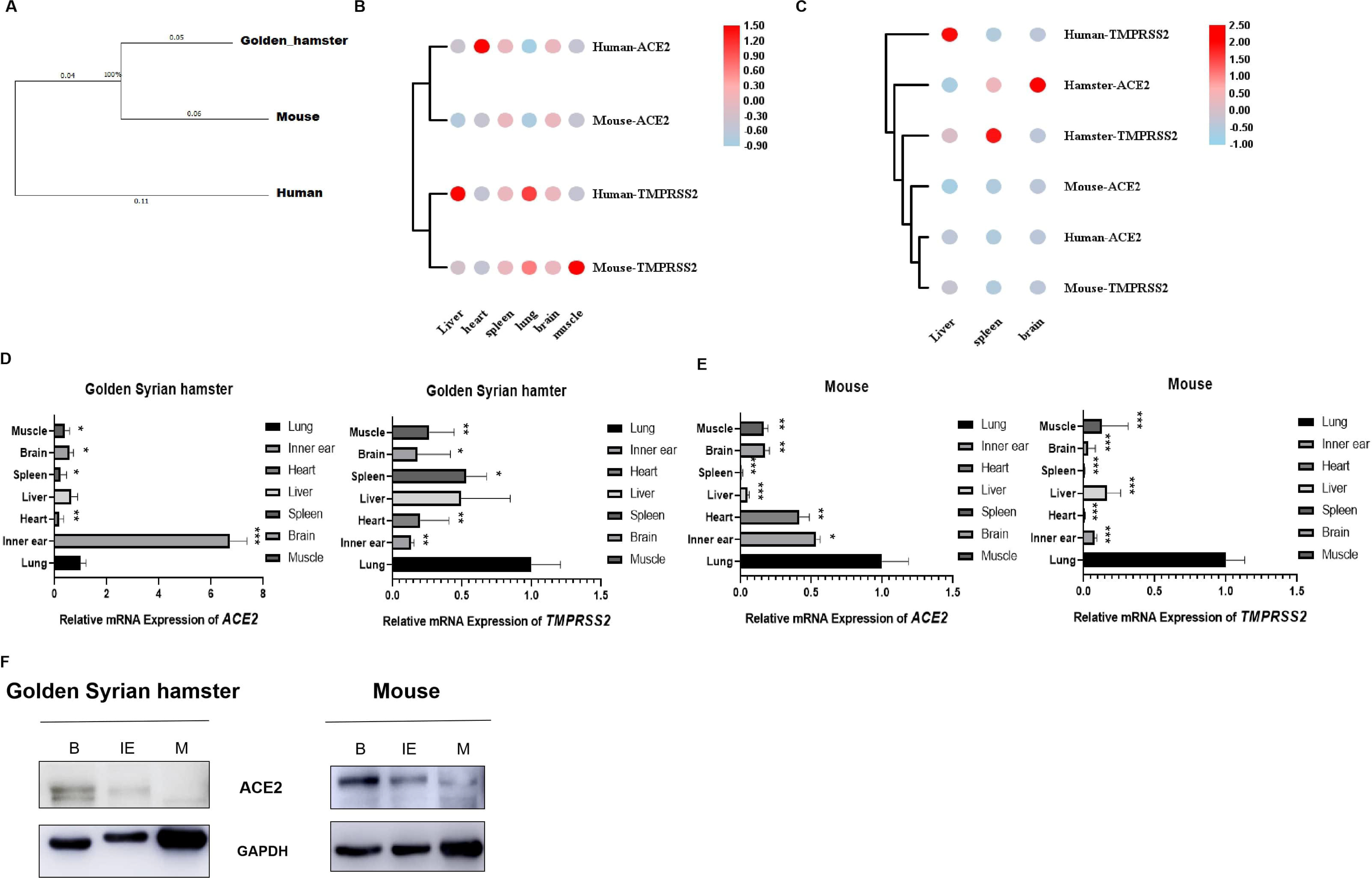
The expression level of *ACE2* and *TMPRSS2* at transcriptomic, DNA, and protein level. A: the phylogenetic of *ACE2* and *TMPRSS2* from human, mouse then the hamster; B: The transcriptome comparison of *ACE2* and *TMPRSS2* from different tissues of human and the mouse; C: The transcriptome comparison of *ACE2* and *TMPRSS2* from different tissues of human, the hamster, and the mouse; D & E: The mRNA expression profile of *ACE2* and *TMPRSS2* in golden Syrian hamster and mouse tissues. Housekeeping gene *Actb* was used for normalization. Relative expression levels (∆∆Ct) of different tissues were compared with that of lung tissue. * *P* <0.05, ** *P* < 0.01, *** *P* < 0.001. F: Expression of ACE2 protein in different tissues of the golden Syrian hamster and mouse. Li: Liver, H: Heart, S: Spleen, L: Lung, B: Brain, IE: Inner ear, M: muscle.

## Discussion

We hypothesized that SARS-CoV-2 might infect the inner ear by binding to *ACE2,* which is the main reason causing sensorineural hearing loss by pathogens invasion. Some studies found *ACE2*, *TMPRSS2,* and *Furin* expressed in the middle ear of a mouse, promising in our data from the present study (Uranaka et al. 2020). Since the transcriptome data from the inner ear specifically has not been available on NCBI, the brain was chosen as an alternative tissue (Zhang et al. 2020). We found *ACE2* did express in the inner ear of both the mouse and the hamster, although the expression level was different compared to *ACE2* expressed in the lung. *TMPRSS2* shows high expression in the lungs compared to it in the inner ear, which might be the reason the symptoms of COVID-19 in the respiratory is more severe in the inner ear. More sophisticated work is required to understand the molecular mechanism of SARS-CoV-2 infecting the inner ear and causing sensorineural hearing loss.

## Author Contribution

Dr. Xue, Dr. Xu, and Dr. Miao designed all the experiments; Dr. Xue did the analysis of transcripts; Dr. Miao and Mrs. Wang finished the qRT-PCR and Western blot; Dr. Tang, Dr. Miao, and Dr. Wang revised the manuscript. All authors are involved in writing and editing this manuscript.

## Conflict of Interest Disclosure

None report

## Funding

This study was supported by the Collaborative Innovation Project of Zhengzhou (Zhengzhou University) (Grant no. 18XTZX12004), the Ph.D. research startup foundation of Henan University of Chinese Medicine (Grant no.RSBSJJ2018-36), Henan Province emergency (Grant no. 201100312300), Zhengzhou emergency project (Grant no. ZZKJ2020604), and the Department of Science and Technology of Henan Province (Grant no. 201100312100).

## Notes

### Competing Interest Statement

The authors have declared no competing interest.

## Reference

Degen C, Lenarz T, Willenborg K. (2020) Acute Profound Sensorineural Hearing Loss After COVID-19 Pneumonia. Mayo Clin Proc 95: 1801–1803.

Harenberg J, Jonas JB, Trecca EMC. (2020) A Liaison between Sudden Sensorineural Hearing Loss and SARS-CoV-2 Infection. Thromb Haemost.

Kilic O, Kalcioglu MT, Cag Y, Tuysuz O, Pektas E, Caskurlu H, Cetin F. (2020) Could sudden sensorineural hearing loss be the sole manifestation of COVID-19? An investigation into SARS-COV-2 in the etiology of sudden sensorineural hearing loss. Int J Infect Dis 97: 208–211.

Langmead B, Salzberg SL. (2012) Fast gapped-read alignment with Bowtie 2. Nat Methods 9: 357–U54.

Uranaka T, Kashio A, Ueha R, Sato T, Bing H, Ying G, Kinoshita M, Kondo K, Yamasoba T. (2020) Expression of Ace2, Tmprss2, and Furin in mouse ear tissue. bioRxiv: 2020.06.23.164335.

Zhang JJ, Dong X, Cao YY, Yuan YD, Yang YB, Yan YQ, Akdis CA, Gao YD. (2020) Clinical characteristics of 140 patients infected with SARS-CoV-2 in Wuhan, China. Allergy.

