## Supplemental methods for "SARS-CoV-2 infecting the inner ear results in potential hearing damage at the early stage or prognosis of COVID-19 in rodents": Supplementary_Materials and Methods.docx

**Animals**

The procedures in this study involving animals were reviewed and approved by the Animal Experimentation Ethics Committee of Henan University of Chinese Medicine. Syrian hamster (n = 3) and C57BL/ 6 mice (n = 6) were sacrificed, and tissues were collected, including inner ear, liver, heart, spleen, lung, kidney, brain and muscle. All tissues were immediately immersed into the TRIzol reagent (TaKaRa, Japan) after dissection and stored at −80℃ before further experiments.

**qRT-PCR**

Total RNA was extracted from tissues homogenates and purified with the TRIzol reagent according to the manufacturer’s protocol. Total RNA (1 μg) was reverse transcribed by PrimerScript RT Reagent Kit (TaKaRa, Japan) following manufacturer instructions. Quantitative real-time reverse transcription polymerase chain reaction (qRT-PCR) reactions were performed using the TB Green Premix Taq II (TaKaRa, Japan) following the manufacturer’s instructions. The genes were amplified by with specific primers (Supplementary Table 1). The reference gene, human β-actin gene and mouse β-actin gene was used for normalization. Sample were run in triplicates and a negative control (no template control) was run concurrently with cDNA to check for primer dimers and contaminants. All reactions were performed twice to ensure technical reproducibly of the assays. The average standard deviation within duplicates studied was 0.5 cycles.

**Western Blotting**

Tissues protein extracts were isolated from inner ear, brain and muscle from Syrian hamster (n = 3) and C57BL/ 6 mice (n = 6) with Tissue Protein Extraction kit (CWBIO, Beijing, China) following manufacturer instructions. Bradford Protein Assay (CWBIO, Beijing, China) was used to estimate the protein concentration. 50 μg of protein supplemented with 5× LSB loading dye was separated in a 10% gel by SDS-PAGE and transferred to Turbo Midi PVDF by semi-dry blotting. After blocking for 1 h at room temperature with a blocking solution of 5 % dry milk, the membrane was incubated with the first antibody diluted in blocking solution at 4 ℃ overnight. Incubation with the secondary, HRP-conjugated antibodies was done for 1 h at room temperature. For detection of the ACE2 and TMPRSS2 protein, a 1:1000 dilution of the ACE2 Rabbit polyclonal antibody (21115-1-AP, Proteintech, USA) and TMPRSS2 (EPR3861) antibody (ab92323, Abcam, USA) was used as the first antibody. And a 1:12000 dilution of a goat anti-Rabbit IgG (HRP) was used as the secondary antibody (ab205719, Abcam, Cambridge, MA, USA). GAPDH expression was demonstrated with a 1:10,000 dilution of GAPDH mouse monoclonal antibody (60004-1-lg, Proteintech, USA) as the first antibody and a 1:12,000 dilution of the goat anti-mouse antibody (ab205719, Abcam, USA) as the second antibody. All data were analyzed with Graphpad Prism 8.0 software (GraphPad Software, United States). Comparison between two groups were performed by Student t-test or unpaired Student t-test. Significance was defined as p < 0.05, data is reported as mean±SEM, and error bars indicate SEM.

**Statistical analysis**

All data were analyzed with Graphpad Prism 8.0 software (GraphPad Software, United States). Comparison between two groups were performed by Student t-test or unpaired Student t-test. Significance was defined as p < 0.05, data is reported as mean±SEM, and error bars indicate SEM.

We found ACE2 and TMPRSS2 co-expressed in inner ear of mouse and hamster as humans, and ACE2 shows high expressed in inner ears compared to lungs while TMPRSS2 shows low expression. Our results indicate there is a potential of SARS-CoV-2 infect inner ear of humans and causing hearing loss at the early stage of COVID-19 and even a profound hearing loss after healing.

**Supplementary Table 1: Primers for *ACE2* and *TMPRSS2* gene expression analysis**

| **Species** | **Genes** | **Primers** |  | **Sequence (5' - 3')** | **Annealing Temp (^o^C)** | **Amplicon size (bp)** | **References** |
| --- | --- | --- | --- | --- | --- | --- | --- |
| Syrian hamster | *ACE2* | Primer 1 | F: | CAATGGTGAATCAGGGCTGG | 60 | 149 | XM_005074209.2 |
|  |  |  | R: | GTGGCAGACCACTTTCCGAT |  |  |  |
|  |  | Primer 2 | F: | ACTGACAATTGTTGGGACGC | 60 | 130 |  |
|  |  |  | R: | ACCAACGATCTCTCGCTTCA |  |  |  |
|  | *TMPRSS2* | Primer 1 | F: | ATCACAGCTCCACAGTTCGC | 60 | 86 | XM_013116227.2 |
|  |  |  | R: | GCACACAGGATACCAGGCTT |  |  |  |
|  |  | Primer 2 | F: | CACTGATCACAGCTCCACAGTTC | 60 | 104 |  |
|  |  |  | R: | TCCAATCATCCTGGCACACAG |  |  |  |
|  | *ACTB* |  | F: | GTGCTATGTTGCCCTGGACT | 60 | 113 | NM_001281595.1 |
|  |  |  | R: | GCTCGTTGCCAATGGTGATG |  |  |  |
| Mouse | *ACE2* | Primer 1 | F: | TCCATTGGTCTTCTGCCATCC | 60 | 198 | Ref |
|  |  |  | R: | AACGATCTCCCGCTTCATCTC |  |  |  |
|  |  | Primer 2 | F: | TGATGAATCAGGGCTGGGATG | 60 | 184 |  |
|  |  |  | R: | ATTCTGAAGTCTCCGTGTCCC |  |  |  |
|  | *TMPRSS2* | Primer 1 | F: | GAGAACCGTTGTGTTCGTCTC | 60 | 180 | ref |
|  |  |  | R: | GCTCTGGTCTGGTATCCCTTG |  |  |  |
|  |  | Primer 2 | F: | AGGATTACAACGCAAGCCTCA | 60 | 177 |  |
|  |  |  | R: | AGAACAGTTGCTGTCCCAGAA |  |  |  |
|  | *ACTB* |  | F: | GATCAAGATCATTGCTCCTCCTGA | 60 | 184 | ref |
|  |  |  | R: | AAGGGTGTAAAACGCAGCTCA |  |  |  |

F: forward primer; R: reverse primer; Temp: temperature; bp: base pairs

Ref：Ma, D., Chen, C., Jhanji, V. et al. Expression of SARS-CoV-2 receptor ACE2 and TMPRSS2 in human primary conjunctival and pterygium cell lines and in mouse cornea. Eye 34, 1212–1219 (2020). https://doi.org/10.1038/s41433-020-0939-4
